# Selective and Improved Photoannealing of Microporous Annealed Particle (MAP) Scaffolds

**DOI:** 10.1101/2020.10.27.358127

**Authors:** Blaise N. Pfaff, Lauren J. Pruett, Nicholas J. Cornell, Joseph de Rutte, Dino Di Carlo, Christopher B. Highley, Donald R. Griffin

## Abstract

Microporous Annealed Particle (MAP) scaffolds consist of a slurry of hydrogel microspheres that undergo annealing to form a solid scaffold. MAP scaffolds have contained functional groups with dual abilities to participate in Michael-type addition (gelation) and radical polymerization (photoannealing). Functional groups with efficient Michael-type additions react with thiols and amines under physiological conditions, limiting usage for therapeutic delivery. We present a heterofunctional maleimide/methacrylamide 4-arm PEG macromer (MethMal) engineered for selective photopolymerization compatible with multiple polymer backbones. Rheology using two classes of photoinitiators demonstrates advantageous photopolymerization capabilities. Functional assays show benefits for therapeutic delivery and 3D printing without impacting cell viability.

## Introduction

Microporous Annealed Particle (MAP) scaffolds have demonstrated great promise as a biomaterial platform in a diverse range of applications from accelerating dermal wound healing to enhancing cell proliferation *in vitro*^1– 4^. MAP scaffolds are composed of a flowable slurry of individual hydrogel microspheres (or microgels) that are transitioned *in situ* to a solid scaffold. This transition occurs through a secondary crosslinking step (i.e. annealing), which chemically bonds the microgels to one another at surface-surface contact points between adjoining microspheres. Multiple annealing chemistries have been demonstrated for MAP scaffolds that vary greatly in mechanism and efficiency (e.g. transglutaminase enzyme reaction, soluble polymeric crosslinkers, and photoinitiated polymerization)^5^. Photoinitiated radical polymerization has particular translational potential due to its speed and compatibility with arthroscopic surgical procedures^6,7^.

Previous iterations of MAP scaffolds that contain maleimide^2^ (Mal) or vinyl sulfone^1^ (VS) groups have dual ability to participate in efficient Michael-type addition necessary for microgel formation as well as undergo radical photopolymerization to anneal (photoanneal). While these functionalities can participate in radical polymerization, this capability cannot be separated from their role in gelation. To allow the incorporation of a functional group that is solely dedicated to photoannealing into any MAP microgels formed via Michael-type addition, we designed a macromer for controlled addition of methacrylamide groups. This approach enables improved translation with its selective radical-initiated activity that is uncomplicated by a parallel ability to participate in undesirable Michael-type reactions.

The controlled release of delivered soluble bioactive molecules (e.g. chemotherapy agents, cytokines) is an important translational function of the hydrogel format approach^8^. However, the presence of nucleophiles (e.g. thiols and primary amines) on many therapeutic molecules severely limits the use of either Mal and VS functional groups for MAP annealing as their ability to participate in Michael-type addition^9^. Specifically, this can promote unintended covalent bonding of their therapeutic cargo to the hydrogel backbone, thus limiting the release of bioactive signals (and, secondarily, impacting availability of functional groups for MAP annealing).

As an alternative to Mal and VS groups, methacrylamide moieties do not participate readily in Michael-type addition under physiological conditions^10^ and, thus, should not limit the ability to incorporate bioactive molecules (e.g. growth factors and drugs). Further, they exhibit a relatively high radical intermediate stability^11^, which we proposed would improve annealing kinetics. To incorporate the methacrylamide functional group for selective photoannealing, we have designed and synthesized a heterofunctional methacrylamide/maleimide (∼3:1) 4-arm poly(ethylene glycol) (PEG) macromer (MethMal). Specifically, the maleimide functional group in MethMal promotes immobilization to the microsphere backbone network during microsphere gelation, while the methacrylamide group is used for free-radical polymerization during photoannealing. The use of methacrylates to anneal MAP scaffolds has been demonstrated previously when methacrylated gelatin (GelMA) was used as both the gelation backbone and the photoannealing agent^12^. However, we believe our presentation of a heterofunctional macromer represents a more robust approach due to a passive retention of the methacrylate groups during microgel formation (i.e. not consumed during this step) and, importantly, an ability to be applied synergistically to a wide array of polymer backbones as a dedicated photoannealing additive.

Here, we report the synthesis, characterization, and use of this custom PEG macromer for annealing MAP scaffolds and provide a comparison of photopolymerization annealing kinetics followed by a series of functional *in vitro* assays, including cell incorporation, protein delivery, and 3D printing.

### Experimental

MethMal was synthesized via a 2-step, one pot modification of 4-arm 20kDa PEG-Maleimide using 2-aminoethanethiol followed by amidation using methacrylic acid via DMTMM (4-(4,6-dimethoxy-1,3,5-triazin-2-yl)-4-methyl-morpholinium chloride). HNMR analysis showed approximately 27% of the arms with a maleimide group and 73% modified with a methacrylamide group (Supplemental Fig. 1) with a 69.4% yield.

To investigate the impact of MethMal on MAP annealing and function, three microgel types with equivalent mechanical properties, particle size, and concentration of annealing functional groups (VS, Mal, or Methacrylamide) were synthesized (Fig. 1A). Using established techniques^5^, we used Instron mechanical testing of macrogels to determine formulations matched with a Young’s modulus of approximately 46kPa (Fig. 1B). Using a previously published microfluidic method^13^ (Fig. 1C), uniform microgels were generated at high throughput with matched microgel diameters (∼80µm) and a low polydispersity index (PDI ≤ 1.05) for all conditions (Fig. 1D-E). All microgel formulations displayed similar post-gelation swelling characteristics (Supplemental Fig. 2), which was important for maintaining the equivalency of annealing groups present following gelation and purification (theoretically 1mM of VS, Mal, or Methacrylamide). Notably, the MethMal microgel formulation used a PEG-Maleimide backbone chemistry and, thus, included a quenching step (via incubation with excess N-Acetyl-L-cysteine) to cap any unreacted maleimides. Importantly, the mechanical and geometric matching achieved through tuning pre-gel formulations and microfluidic generation, respectively, allowed us to isolate the photoannealing chemistry (MethMal, Mal, or VS) as the only variable for subsequent studies.

**Figure 1.**
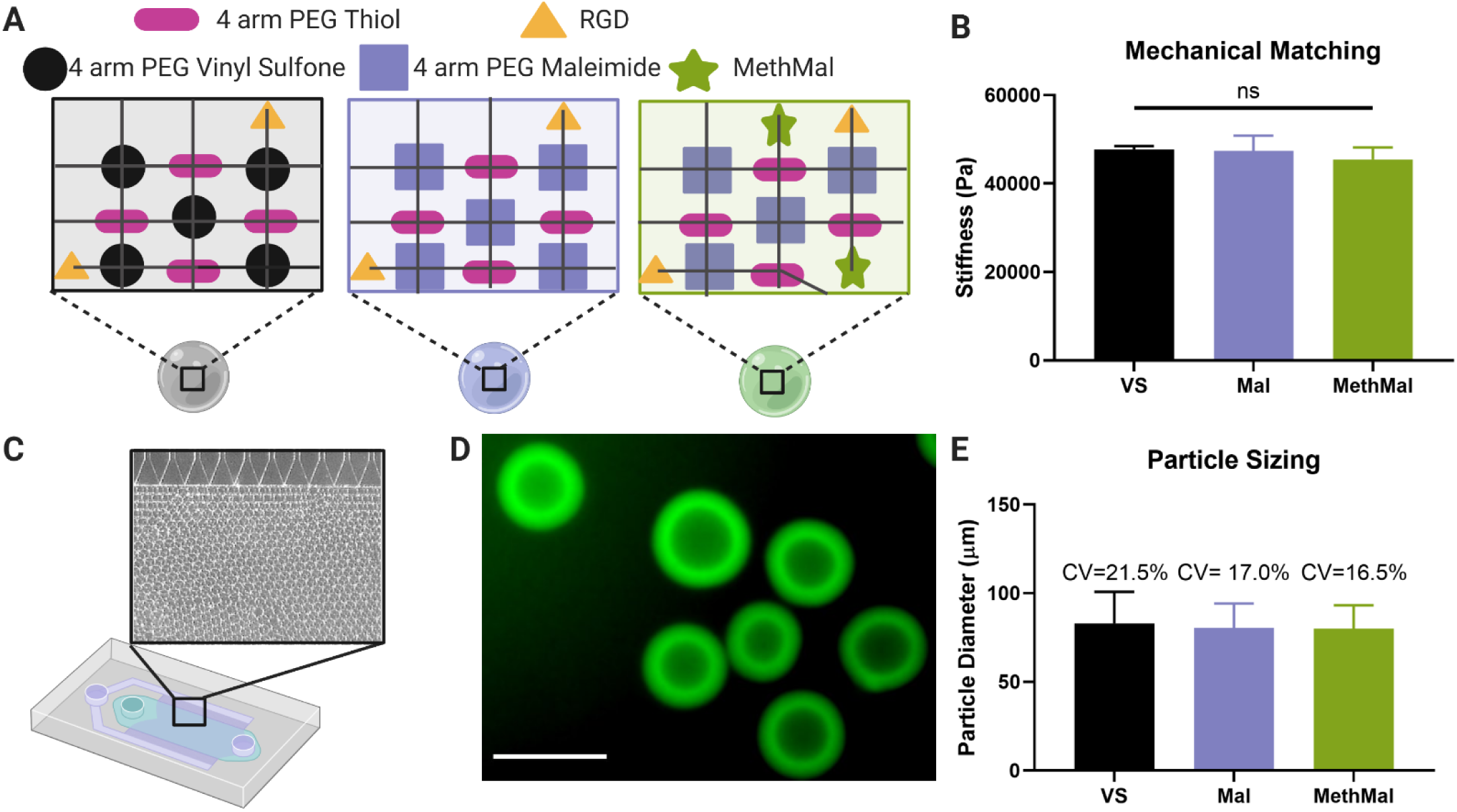
Synthesis and characterization of three microgel types. A) The three gel formulations were composed of a PEG-Vinyl Sulfone backbone (VS), a PEG-Maleimide backbone (Mal), and a PEG-Maleimide with 1mM MethMal (MethMal). All gel formulations were crosslinked with a 4-arm PEG Thiol and had RGD cell adhesive peptide. B) The three gel formulations were mechanically matched to have a Young’s modulus of ∼46kPa determined by Instron testing of macroscale gels. Note: All microgel compositions were formulated to stoichiometrically provide a theoretical 1mM excess photoannealing functional group for all conditions. C) Microgels were produced using a high throughput microfluidics device. D) Biotinylated microgels were fluorescently visualized with streptavidin-488. Scale bar represents 100µm. E) Microgels were size matched to approximately ∼75µm in diameter. All graphs show mean +/-standard deviation. One-way ANOVA was used to determine significance in mechanical moduli.

Our first comparative study involved rheological analysis to observe photoannealing kinetics using multiple photoinitiators relevant to biologic applications. Specifically, we chose to focus on lithium phenyl-2,4,6-trimethylbenzoylphosphinate (LAP) as a type 1 photoinitiator due to its common usage for hydrogel formation and Eosin-Y as a type 2 photoinitiator due to the ability to penetrate deeper into tissues with visible wavelengths^6,14^. Interestingly, for all conditions exposed to 1mM LAP a noticeable decrease in storage modulus was observed after extended light exposure that is indicative of an unintended degradation event (Fig. 2A) during radical activation^15^. All scaffold conditions demonstrated logarithmic annealing behavior for each photoinitiator (Fig. 2A) demonstrating each functional group is capable of undergoing radical polymerization necessary for annealing. For a given amount of energy, MethMal produced a significantly higher change in storage modulus than Mal using 0.1mM LAP and a significantly higher change than VS using Eosin-Y (i.e. demonstrating greater annealing efficiency, Fig. 2B). Additionally, MethMal required significantly less energy to reach 50% of the maximum storage modulus than VS and Mal using 0.1mM LAP and less than VS using Eosin-Y (Fig. 2C). We observed noticeable differences between LAP and Eosin-Y annealing behavior which we attribute to their identity as type 1 and type 2 photoinitators, and we plan to investigate this in future studies.

**Figure 2.**
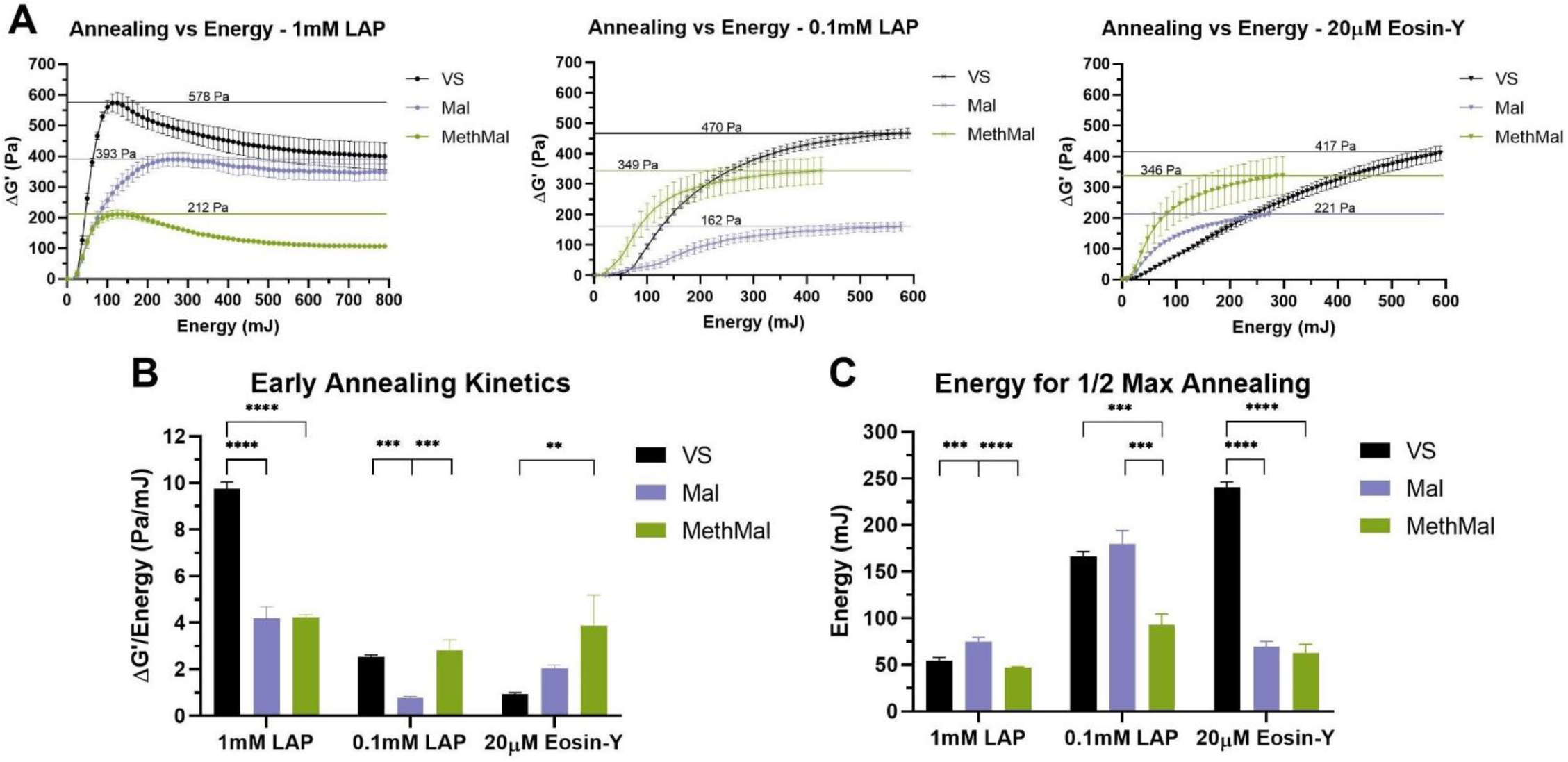
Quantification of annealing across photoinitiators via rheological analysis. A) Change in storage moduli compared to light energy introduced to the system. Horizontal lines indicate maximum ΔG’. B) Early annealing kinetics determined by maximum rates of change following toe regions of curve. C) Light energy required to reach one-half of the maximum increases in storage moduli. All graphs show mean +/-standard deviation. One-way ANOVAs followed by post-hoc multiple comparisons tests (Tukey HSD) were used to determine significance. **** p-value < 0.0001, *** p < 0.001, ** p < 0.01.

All annealing chemistries and photoinitiator conditions were compared in a cell viability assay using primary adult human dermal fibroblasts (HDFs). Briefly, cells were mixed with unannealed MAP microgels before undergoing annealing. Each condition was exposed to light for the amount of time required to achieve maximum annealing (Supplemental Table 1). Cell viability was determined at 24 hours following the annealing step (Fig. 3A). MethMal consistently demonstrated the highest average cell viability among all photoinitiator conditions (Fig. 3B). Mal gels showed significantly lower viability than MethMal and VS gels during 1mM LAP annealing. In addition, VS gels displayed statistically significantly lower viability than MethMal for 0.1mM LAP annealing. All chemistries exhibited reduced viability during Eosin-Y annealing. This may be a result of the longer light exposure times required to reach max annealing with Eosin-Y. To further investigate this possibility, we repeated the MAP viability assay with HDFs seeded in MethMal gel and varied the time of exposure used for annealing. We observed a clear reduction in cell viability as annealing time increased (Fig. 3C). These results emphasized the importance of reducing the time of radical exposure that is provided by MethMal (e.g. reduced energy to reach 50% annealing, Fig. 2C).

**Figure 3.**
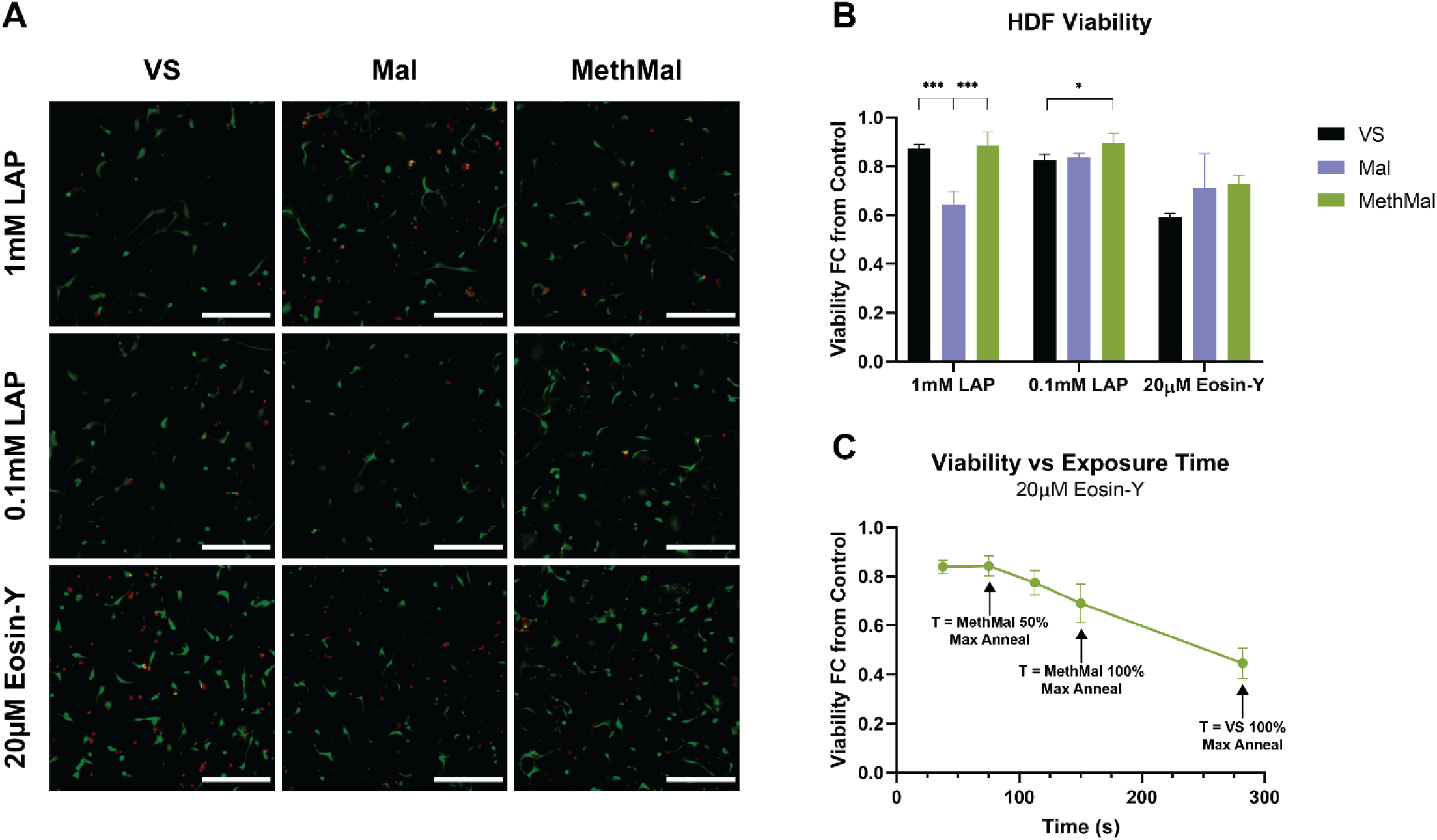
Primary cell viability in response to annealing chemistries and photoinitiators. A) Representative maximum intensity projection (MIP) images from each condition showing both live (green) and dead (red) HDFs. Scale bar represents 300µm. B) HDF viability at 24 hours shown as a fold-change from 2D tissue culture plastic controls. C) HDF viability at 24 hours following culture in MethMal gel and annealed with 20µM Eosin-Y using a range of exposure times. All graphs show mean +/-standard deviation. One-way ANOVAs followed by post-hoc multiple comparison tests (Tukey HSD) were used to determine significance. *** p-value < 0.001, * p < 0.05.

To investigate the functional benefits of the MethMal annealing macromer, we explored two current biomaterial applications: therapeutic delivery and 3D printing. Therapeutic delivery often requires release of molecules that contain free thiol groups (e.g. proteins) which can readily interact with either the VS or Mal groups prior to annealing^9^. To determine the differences in protein release from the three gel types, as well as the importance of fully processing (i.e. quenching the leftover maleimide groups) the MethMal condition, fluorescently tagged bovine serum albumin (BSA) release was quantified for 72 hours under infinite sink conditions (Fig. 4A). Starting at 24 hours, the quenched MethMal group demonstrated significantly more protein release than all other groups as determined by fluorescence intensity in the supernatant. By 72 hours, the release profiles had plateaued and the quenched MethMal group had released all of the loaded BSA (Fig. 4B). Notably, the other groups had not fully released the BSA and in particular the Mal gel only released approximately 50%. In a separate investigation, we found that exposing the Mal and VS conditions to excess thiol-containing molecules resulted in a clear loss of annealing capacity (Supplemental Fig. 6). Overall, this model protein release assay demonstrated the impact of using a dedicated annealing chemistry (i.e. methacrylamide) on the ability of MAP to act as a therapeutic delivery depot.

**Figure 4.**
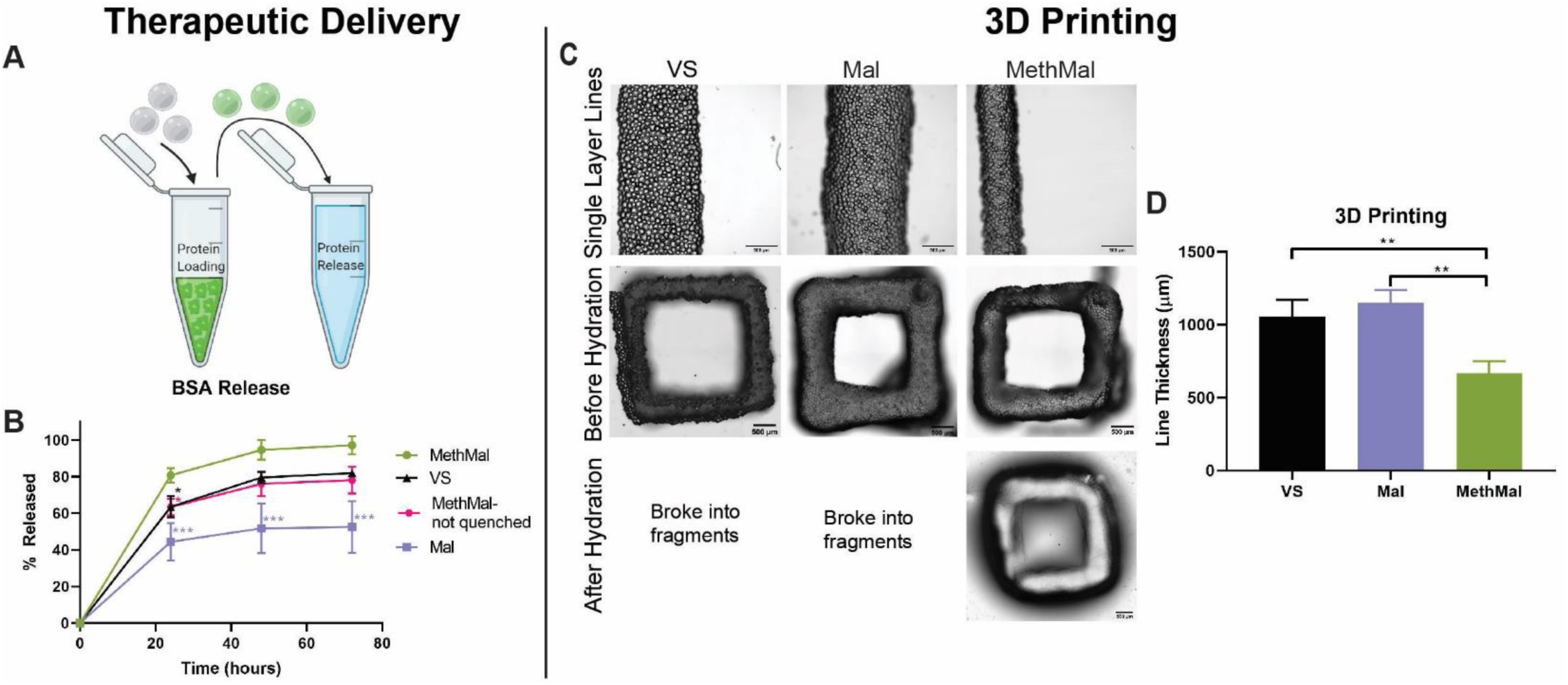
Functional assays to compare annealing chemistries. A) BSA release was characterized by loading MAP particles with fluorescently tagged BSA and monitoring release in infinite sink conditions for 72 hours. B) Release profiles over 72 hours. C) 3D printing experiments consisted of printing single-layer lines and 5-layer squares and submerging them in PBS to test the quality of crosslinking. D) Line thickness was compared between groups to evaluate the kinetics of annealing. Scale bars represent 500µm. All graphs show mean +/-standard deviation. One-way ANOVAs followed by post-hoc multiple comparisons tests (Tukey HSD) were used to determine significance. *** p-value < 0.001, ** p < 0.01, * p < 0.05.

For the purposes of 3D printing, rapid covalent stabilization of printed structures is necessary to maintain shape fidelity of physically stabilized filaments to match computer designs^16–18^. To compare the impact of photoannealing functional group choice on extrusion-based 3D printing, we used two proof of concept experiments with Eosin-Y as a photoinitiator. In the first experiment, lines were printed at a uniform translation speed (100mm/min). After printing, the gels were imaged and analyzed for the average line thickness to determine the precision of printing (Fig. 4D). The MethMal group had significantly thinner lines which can be contributed to the quicker annealing kinetics for the MethMal group, which prevented settling and flattening of the filament^18^ that occurred in the other groups immediately upon deposition (Fig. 2B). In the second experiment, 5-layer tall squares were printed and submerged in PBS for 5 minutes after printing to determine the functional stability of annealing. After 5 minutes, only the MethMal group remained intact (Fig. 4C), while the Mal and VS gels had broken into small fragments (Supplemental Fig. 8), indicating that MethMal annealing provided both rapid and more stable annealing within filaments upon deposition. Notably, these two 3D printing experiments were conducted using standard protocols and set-ups for non-MAP based inks and, therefore, we believe that the print resolution (∼600µm) can be greatly enhanced with targeted device and protocol changes, which we will explore in future studies. In summary, these proof of concept experiments provide clear evidence that the MethMal chemistry provides printing resolution and stability advantages for 3D printing of MAP scaffolds.

## Conclusions

In conclusion, we synthesized a targeted heterogeneous macromer, MethMal, for the selective purpose of enhancing MAP photoannealing capabilities. MethMal can easily be incorporated into MAP gels of any formulation, provided the microgel gelation utilizes a Michael-type addition mechanism. Further, MethMal displayed enhanced photoannealing without impacting cell viability. These advantages in annealing kinetics, combined with reactive selectivity (i.e. no unintended immobilization of nucleophile-containing molecules), translate into the MethMal chemistry providing clear advantages in the areas of therapeutic delivery and 3D printing as shown by our proof of concept experiments.

## Acknowledgements

Anton Paar Modular Compact Rheometer and OmniCure Series 2000 located in the Caliari Lab (Center for Advanced Biomanufacturing). Figure 1 and 4 schematics created with BioRender.com. LP was supported by a National Science Foundation Graduate Research Fellowship and by the National Heart, Lung, and Blood Institute of the National Institutes of Health under Award Number F31HL154731. BP and NC were supported by the National Institutes of Health Biotechnology Training Program (T32GM008715). This work was partially supported through the US National Institutes of Health High Priority, Short-Term Project Award (1R56DK126020-01) and The Wallace H. Coulter Translational Partners Program at The University of Virginia.

