## Supplemental Information for "Selective and Improved Photoannealing of Microporous Annealed Particle (MAP) Scaffolds"

**Supplementary Information:**

***Methods***

**Biomaterial Creation and Characterization:**

*Sources of Materials:* Four-arm poly(ethylene glycol) maleimide (PEG-Mal, 10kDa and 20kDa) and poly(ethylene glycol) thiol (PEG-SH, 10kDa) were purchased from Nippon Oil Foundry (Japan). Poly(ethylene glycol) vinyl sulfone (PEG-VS, 10kDa) was purchased from Jenkem Technology (USA). RGD cell adhesive peptide (Ac-RGDSPGGC-$NH_2$) was purchased from WatsonBio Sciences (USA). All materials were dissolved into either ultrapure water or 0.1% trifluoroacetic acid solutions (to prevent disulfide bond formation) and aliquoted into specific amounts to ensure precision. The aliquots were lyophilized and stored at -20°C until use. LAP was purchased from Sigma-Aldrich (USA) and Eosin-Y from Acros Organics (Belgium).

*Heterofunctional maleimide-methacrylamide 4-arm PEG Macromer Synthesis:* Four-arm poly(ethylene glycol) maleimide (20kDa) was dissolved in pH=7.4 1X PBS for 30min at room temperature at a concentration of 1g per 10mL. 2-aminoethanethiol (Acros Organics) was added at a 0.67:1 thiol-to-maleimide molar ratio and reacted for 2 hours at room temperature. DMTMM (Oakwood Chemical) at a 12:1 molar ratio to maleimides and methacrylic acid (Sigma) at a 8:1 molar ratio of maleimides were pre-reacted for 50 minutes in pH=7.4 1X PBS before being added to the reaction. Triethylamine (Fisher) was added as a catalytic reagent at the same molar ratio as the methacrylic acid and the reaction proceeded overnight at room temperature. The solution was dialyzed (3.5kDa Snakeskin tubing, Fisher) for 3 days in 4L of 1M NaCl (changed 2x daily) followed by distilled water (4L, 1hr x 6 washes). The product was frozen and lyophilized, then analyzed using HNMR.

*Annealing Macromer Characterization:* A Varian Inova 500 NMR spectrometer located in the UVA Biomolecular Magnetic Resonance Facility was used to acquire the spectra. 25mg of the reaction was dissolved in *d*-chloroform (Sigma). MestReNova was used for analysis, and maleimide peak (~6.7 ppm) was compared to the methacrylamide peaks (~5.75 and 5.25 ppm) to determine relative functional group ratio (Supplemental Fig. 1).

*MethMal Pre-Gel Solution Formulation:* The MethMal gel was a 5wt% gel with a PEG-Maleimide backbone. To prepare PEG-Mal, MethMal and RGD was dissolved in pH=4 10X PBS for macrogel production or pH=2 10X PBS for microgel synthesis. This was combined with a PEG-SH crosslinker solution with 5µm biotin-maleimide which was dissolved in ultrapure water. The final concentrations in the gel solution were 32.70mg/mL PEG-Mal, 25.15mg/mL PEG-SH, 0.82mg/mL RGD, and 8.06mg/mL of MethMal.

*Maleimide Pre-Gel Solution Formulation:* The maleimide gel was a 5.3wt% gel with a PEG-Maleimide backbone. To prepare PEG-Mal and 1mM RGD was dissolved in pH=4 10X PBS for macrogel production or pH=2 10X PBS for microgel synthesis. This was combined with a PEG-SH crosslinker solution with 5µm biotin-maleimide which was dissolved in ultrapure water. The final concentrations in the gel solution were 34.38mg/mL PEG-Mal, 25.19mg/mL PEG-SH, and 0.82mg/mL of RGD.

*Vinyl Sulfone Pre-Gel Solution Formulation:* The vinyl sulfone gel was a 5.8wt% gel with a PEG-Vinyl Sulfone backbone. To prepare PEG-VS and 1mM RGD was dissolved in pH=7.3 a 0.3mM triethanolamine buffer for macrogel and microgel production. This was combined with a PEG-SH crosslinker solution with 5µm biotin-maleimide which was dissolved in ultrapure water. The final concentrations in the gel solution were 36.44mg/mL PEG-VS, 31.71mg/mL PEG-SH, and 0.82mg/mL of RGD.

*Macrogel Synthesis:* 150µL gels (i.e. macrogels) were used to mechanically match hydrogel stiffnesses between the three groups to ~45kPa. 150µL of pre-gel solution were formed into pucks between two SigmaCote coated glass microscope slides for thirty minutes past full gelation, which was determined simultaneously using a HAAKE Rheowin viscometer to monitor a plateau in storage modulus. The macrogels were weighed following gelation and then swollen to equilibrium in PBS overnight at 37°C and weighed again to determine the equilibration ratio (Supplemental Fig. 3) prior to mechanical testing.

*Mechanical Analysis of Macrogels:* Macrogels were removed from PBS and excess moisture was wicked with a KimWipe prior to testing, with three macrogels being tested per gel formulation. An Instron mechanical load device was used to test compressive stiffness of the macrogel at a rate of 0.5mm/min for 1mm. Bluehill software analyzed the load (N) and extension (mm) and stress-strain curves were produced. MATLAB shape language modeling software was used to calculate the Young's modulus.

*Microgel Production:* Microgels were produced using a microfluidics PDMS mold created using previously published design created by Rutte et al.[13] in a dust free hood. Briefly, a 1% Pico-Surf surfactant (Sphere Fluidics) solution diluted in NOVEC 7500 oil (3M) was run through the oil channel and the gel formulations described above were in the aqueous channel. Using a syringe pump, the surfactant and gel solutions were run at 5mL/hr in the device and collected in a conical tube. The two microgel formulations (MethMal and Mal only) consisting of a PEG-Maleimide backbone were mixed with a triethylamine solution (20µL TEA/ mL of gel) to increase the pH and accelerate gelation. The VS formulation was not purified until complete gelation was observed as measured using the viscometer (Supplemental Fig. 3) as described above for macrogel synthesis.

*Microgel Purification:* Microgels were washed with NOVEC 7500 oil three times (1X volume of gel). Next, microgels were combined with PBS (5X volume of gel) and washed with NOVEC 7500 oil (1X volume of gel) three times, allowing separation of the oil and aqueous solution by settling. Finally, the oil was removed and the microgels were washed with PBS (3X volume of gel) and hexanes (3X volume of gel) three times. After the final wash, microgels with MethMal reacted overnight at 37°C with a 100mM N-Acetyl-L-Cysteine solution in 1X PBS to cap any excess maleimides.

*Microgel Sterilization:* Microgels were removed from solution via centrifugation (4696gx5min) and all further steps were performed in a biosafety cabinet. Microgels were washed three times with 70% isopropyl alcohol followed by four washes with sterile 1X PBS (pH=7.4) before being mixed with a photoinitiator solution.

**Rheological Analysis of Annealing Properties:**

*Preparation:* MAP particles were isolated via sequential centrifugation steps (4696gx5min followed by 18000gx5min) and aspiration of supernatant. Particles were mixed at a 1:1 ratio with photoinitiator solutions at 2x desired final concentrations in pH=7.4 1X PBS (i.e. LAP 2mM or 0.2mM and Eosin-Y 40µM). After overnight incubation, particles were isolated from excess solution via sequential centrifugation (17000gx5min twice) and aspiration of supernatant. All isolated particles were dried (i.e. removal of liquid between particles) to achieve a similar initial storage modulus starting range (2500 ± 250Pa) prior to annealing on the Anton Paar Modular Compact Rheometer (Supplemental Fig. 4). In cases where the initial storage modulus was outside this specified range (indicating a difference in particle dryness), excess volume was adjusted. Specifically, excess supernatant was either added or removed via re-introduction of supernatant or manual "wicking" using a Kimwipe (Kimtech), respectively.

*Rheology:* An Anton Paar Modular Compact Rheometer (MCR 302) was used to acquire data. Dried particle mixtures (25µL) were placed on the center of the UV stage and an 8mm parallel plate probe (Anton Paar, Part No. 97676) was lowered to 1mm above the sample using Anton Paar RheoCompass software. A custom oscillatory time sweep (1% strain, 1rad/s angular frequency)[19] was performed at 37°C using an initial 0.25N normal force and recording the storage modulus (G') every 0.5s (N=3 per condition). After 5 minutes, the light source (OmniCure Series 2000) was turned on, beginning the annealing process (365nm at 8.35mW/cm2 for LAP, or 400-500nm at 4.33mW/cm2 for Eosin-Y).

*Analysis:* Storage modulus data points were plotted against time in Microsoft Excel. Time was converted to energy to compare across photoinitiators using the equation Energy = time x intensity x cross-sectional area. Data were fit to polynomial curves ($R^2$>0.99) and plugged into a custom MATLAB code to determine inflection points. Inflection points, in conjunction with changes to slope, were used to separate photoannealing properties from natural drying behavior of hydrogels. Early annealing kinetics were determined via analysis of slope following initial toe region of curves (Supplemental Fig. 5).

**Cellular Studies:**

*Source of Cells:* Adult primary human dermal fibroblasts (HDFs) were purchased from ATCC and expanded to passage 5 (P5) via manufacturer's protocol. Briefly, cells were plated on T-75 flasks (ThermoFisher) at a density of 2500-5000 cells/cm$^2$, cultured in complete growth media (ATCC Fibroblast Basal Media and Fibroblast Growth Kit-Low Serum), and passaged at 80-100% confluence. Upon collection at P5, cells were transferred to freezing media (complete growth media with 10% dimethyl sulfoxide) and cryopreserved in liquid nitrogen.

*Cell Viability Assay:* HDFs at P5 were thawed from cryopreservation and expanded on a T-75 flask for 48 hours, with complete growth media being exchanged after 24 hours. At 48 hours cells were passaged and seeded at 5000 cells/cm$^2$ on 56.7cm$^2$ petri dishes (Falcon). Cells reached 80% confluence prior to experimentation. Sterile MAP particles microgels were dried through successive 5-minute centrifugation steps each followed by supernatant aspiration (4696g, 18000g, 25000g) to remove excess volume. Dried gels were mixed at a 1:1 ratio with one 2X photoinitiator of (0.2mM LAP, 2mM LAP, or 40µM Eosin-Y) dissolved in sterile PBS. Following addition of the photoinitiator solution, microgels were incubated at room temperature for 15 minutes before being dried at 25000g for 5 minutes. Cells were collected, counted using a hemocytometer, and pelleted at 500g for 5 minutes. Using a positive displacement pipette, dry microgels MAP particles were used to resuspend the cell pellet at a density of 1000 cells/µL. Individual scaffolds were formed by pipetting 15µL of cell/gel mixture into a 48-well plate (Falcon). The pipette tip was used to spread the gel into an even, thin layer across the bottom of the well. Scaffolds were annealed individually based on gel type and photoinitiator as described in Table S1. Annealing was done for LAP and Eosin-Y conditions at an irradiance of 8.35mW/cm$^2$ and 4.33mW/cm$^2$ respectively, measured immediately before experimentation (ThorLabs PM100D). N=4 scaffolds were created for each gel and each annealing condition. Following annealing, 500µL complete growth media was added to each scaffold well, and the scaffolds incubated at 37°C for 24 hours. For 2D controls, HDFs from the same plate were pelleted and resuspended at a density of 1000 cells/µL in complete growth media. 30µL of cells were plated onto a 24-well tissue culture treated plate (Costar) and 1mL complete growth media was added before incubating at 37°C for 24 hours. Scaffolds and controls were washed once with sterile PBS then stained using a LIVE/DEAD staining kit (ThermoFisher) based on manufacturer instructions. Scaffolds were imaged using both the FITC and Texas Red filters of an ImageXpress Micro Confocal. Briefly, a 50µm z-stack with 5µm steps was centered at the approximate midpoint of the scaffold and compiled into a Maximum Intensity Projection (MIP). Exposure values for both channels were set at 500ms for all scaffolds. N=9 sites were acquired for each macro scaffold to yield live and dead cell counts for each scaffold. Cells were automatically counted quantified via the ImageXpress "Auto-Segment" module followed by a minimum area filter of 50 pixels to minimize artifacts. Control conditions were imaged using the same objective and exposure settings but as 2D images rather than z-series. Automatic quantification for control cells was done using the ImageXpress "Find Blobs" module (Approximate Minimum Width of 5µm, Approximate Maximum Width of 30µm, and Intensity Above Local Background of 500) followed by a minimum area filter of 50 pixels to minimize artifacts. Cell viability for each scaffold was calculated as a percentage of live cells divided by total cells then converted to a fold change above controls by dividing by average control viability.

**Protein Studies:**

*Fluorescent tagging of BSA:* Bovine serum albumin (Fisher Scientific) was dissolved at 5mg/mL in PBS. Alexa Fluor 488 carboxylic acid succinimidyl ester (Invitrogen) was dissolved in dimethylformamide (Sigma) at 2mg/mL. 125µL of reactive dye was added to react for 1 hour at room temperature while shaking. Following the reaction, the unreacted dye was removed using Zeba Spin Desalting columns (Thermofisher Scientific) and the labeled BSA was stored at 4°C until further use.

*Protein Loading MAP gel:* 50µL of MAP gel was resuspended at a one-to-one ratio with a 100g/mL protein solution and incubated for 48 hours while rotating at 4°C. The solutions were then filtered through a 0.22µm filter to separate the gel from the supernatant solution. The supernatant solutions were saved to analyze the loading efficiency and determine the final concentration in the gel.

*Protein Release from MAP gel:* 50µl of MAP gel loaded with BSA was mixed in 20X volume of PBS for infinite sink release conditions. Every 24 hours low speed centrifugation (5minx500g) was used to separate the gel from the solution. The supernatant was saved and new PBS was added to the gel to obtain release profiles. Release was continued for 72 hours. After 72 hours the supernatant solutions (N=3) were analyzed by reading the fluorescence intensity on a plate reader and extrapolating the values with a standard curve of fluorescent BSA.

**3D Printing Studies**

*Gel Preparation:* Microgels were mixed one to one with a 40µM Eosin-Y solution and stored at 4°C overnight. Microgels were dried by going through two high speed centrifugations (17000gx5min) and pouring off the excess photo initiator solution. Dried microgels were loaded into 1mL syringes (Air-Tite) with a 25-gauge needle.

*3D Printer:* A FELIX 3.1 extrusion-based 3D printer from FELIXprinters (IJsselstein, Netherlands) was modified to extrude from a syringe, using a previously published design[20] (Supplemental Fig. 7). Repetier-Host software (freely available from Repetier) was used to control extrusion. A translation speed of 100mm/min was used for printing which correlated to an approximate volumetric extrusion rate of 69µL/min.

*3D Printing Line Thickness:* Repetier- Host software was used to extrude 3 lines for each condition of at least 5mm in length. A ThorLabs 505nm LED (M505L3) was placed to shine on the stage and was turned on for the duration of printing at full intensity (1000mA). After printing the lines of particles were imaged with a Leica DMi8 widefield microscope. 3 lines were printed for each condition and the average thickness was quantified using ImageJ to measure the area of the line divided by the length of the line.

*3D Printing Squares:* Repetier-Host software was used to extrude 5-layer thickness squares. A ThorLabs 505nm LED was placed to shine on the stage and turned on for the duration of printing at full intensity (1000mA). After printing the squares were imaged on a Leica microscope and then placed in a dish of PBS for 5 minutes to test the stability of crosslinking. After 5 minutes the squares were imaged again if they had not broken apart.

***Supplemental Figures:***

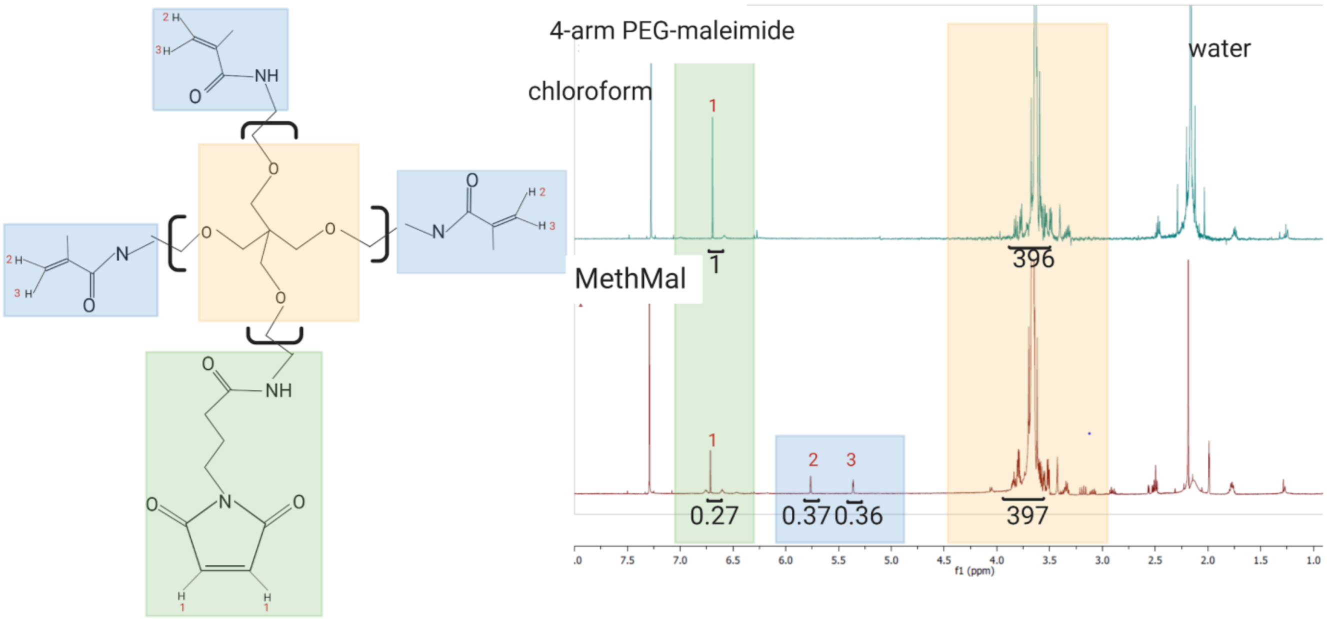


*Supplemental Figure 1:* The MethMal annealing macromer was composed of 20kDa 4-arm poly(ethylene glycol) modified with ~3 methacrylamide (ppm 5.36, 5.76) arms and ~1 maleimide (ppm 6.71) arm.

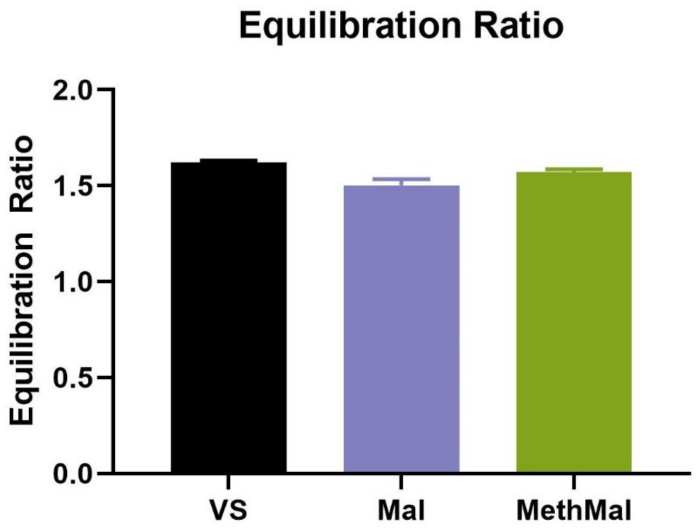


*Supplemental Figure 2:* Equilibration (i.e. change in gel volume following placement in PBS after gelation) ratios for each gel condition.

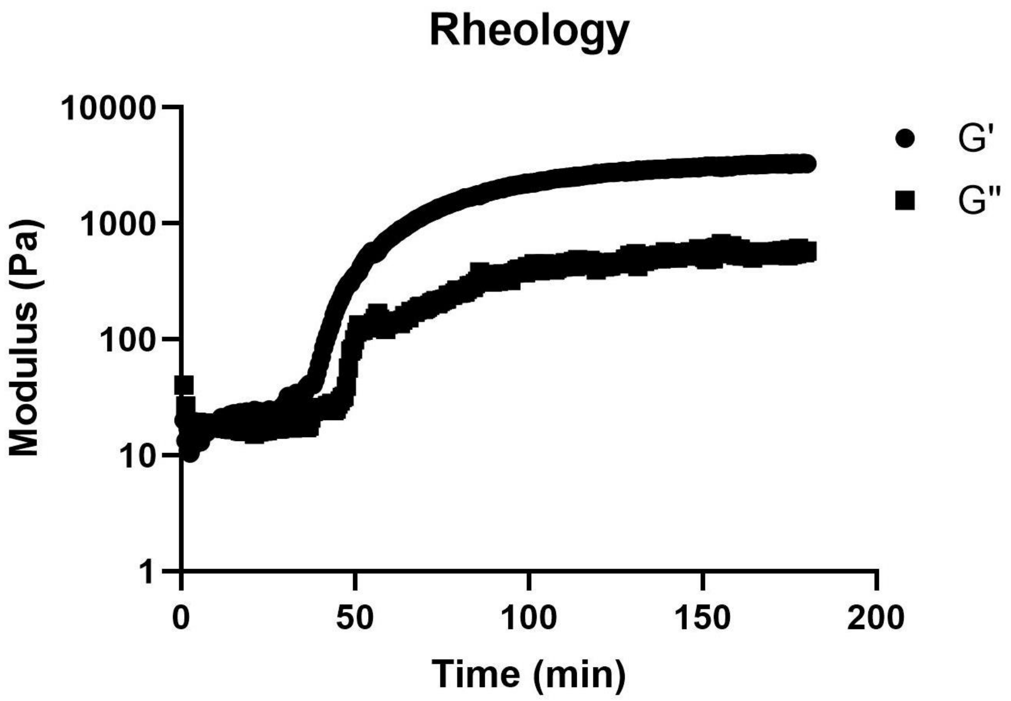


*Supplemental Figure 3:* Rheology for the PEG-VS gel formulation showed gelation started after 41 minutes.

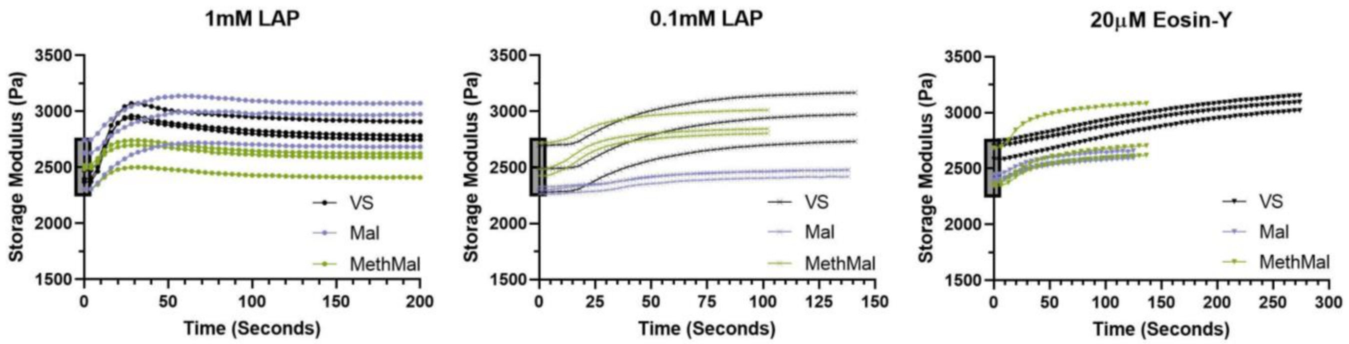


*Supplemental Figure 4:* Raw rheology data across photoinitiators for quenched MethMal and unquenched Vinyl Sulfone and Maleimide gels. Demonstrates that all storage moduli initial values fell between 2500 +/- 250Pa.

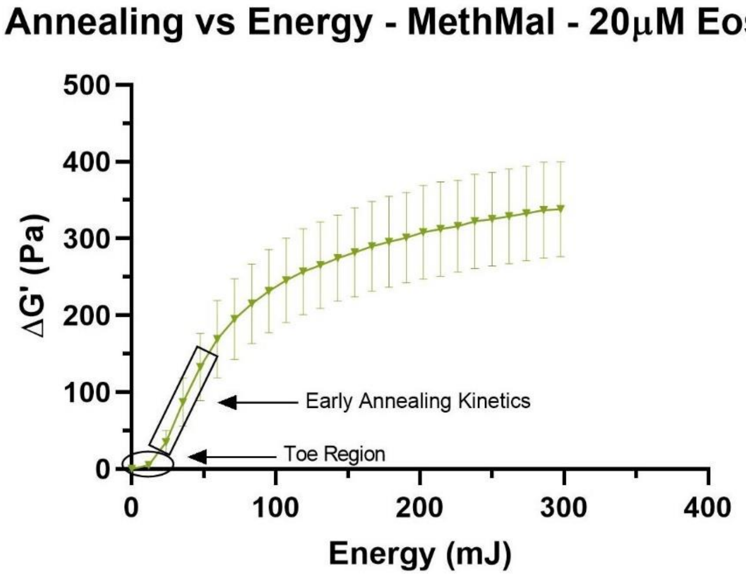


*Supplemental Figure 5:* Example analysis of early annealing kinetics. Maximum slopes were recorded following the initial toe region of the curve.

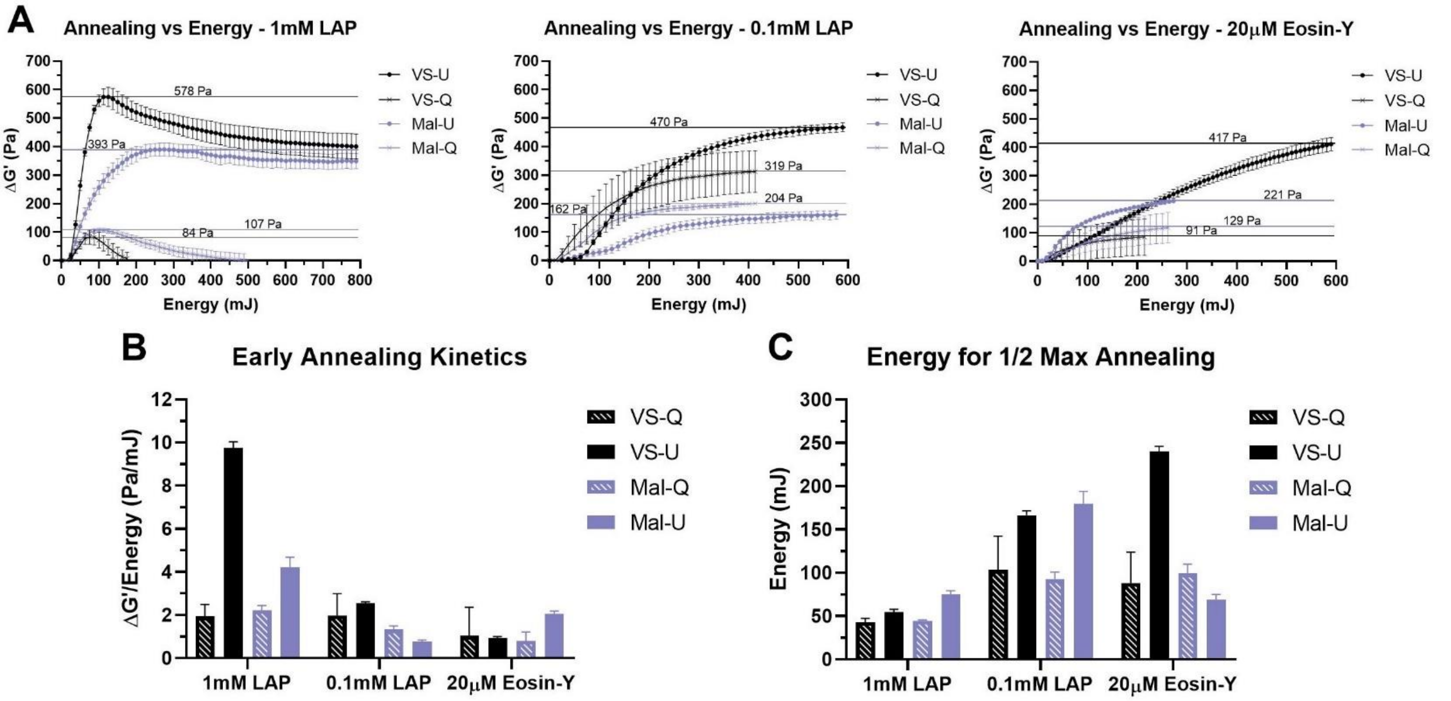


*Supplemental Figure 6:* Quantification of annealing of quenched Vinyl Sulfone (VS-Q) and Maleimide (Mal-Q) gels as well as unquenched Vinyl Sulfone (VS-U) and Maleimide (Mal-U) gels from Figure 2 across photoinitiators via rheological analysis. A) Change in storage moduli compared to light energy introduced to the system. Horizontal lines indicate maximum ΔG’. B) Early annealing kinetics determined by maximum rates of change following toe portions of curve. C) Light energy required to reach one-half of the maximum increases in storage moduli. All graphs show mean +/- standard deviation.

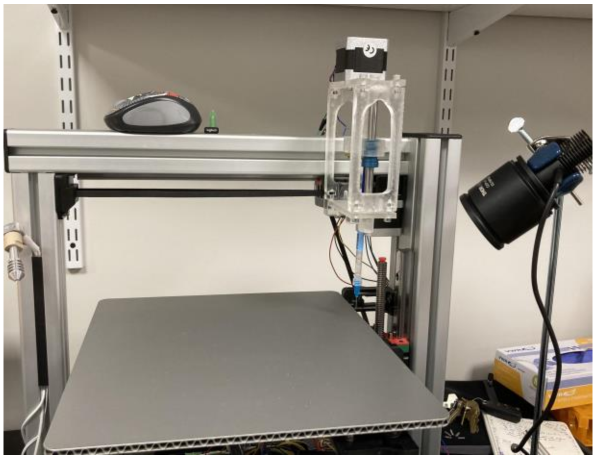

*Supplemental Figure 7:* 3D printing setup.

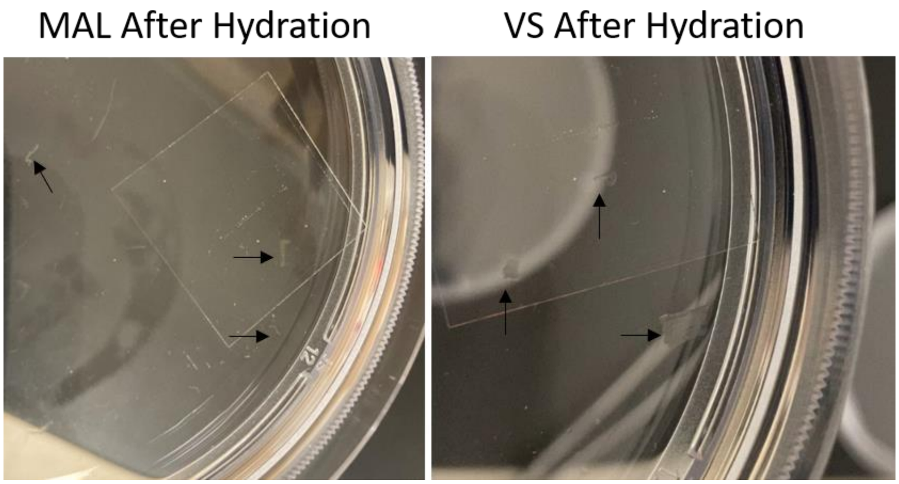


*Supplemental Figure 8:* 3D printed squares of MAL and VS gels broke into fragments after being submerged in PBS. Arrows point to gel fragments in the petri dish.

**Table S1:** Time (seconds) to max annealing based on photoinitiator and gel type.

| | **VS** | **MethMal** | **Mal** |
|---|---|---|---|
| **1mM LAP** | 28.3 | 30.7 | 70.8 |
| **0.1mM LAP** | 145.0 | 113.8 | 141.0 |
| **20μM Eosin-Y** | 282.1 | 150.0 | 147.4 |
